# Altered defense patterns upon retrotransposition highlights the potential for rapid adaptation by transposable elements

**DOI:** 10.1101/2023.12.20.572632

**Authors:** Emma S.T. Aller, Christa Kanstrup, Pascal Hunziker, Daniel J. Kliebenstein, Meike Burow

## Abstract

Transposable elements can be activated in response to environmental changes and lead to changes in DNA sequence. Their target sites of insertions have previously been thought to be random, but this theory has lately been contradicted. For instance, mobilization is favored towards genes involved in regulatory processes. This makes them interesting as potential players in rapid responses required under stressful environmental conditions. In this paper, we report the in-depth characterization of an *Arabidopsis thaliana* Col-0-based line whose altered DNA methylation pattern made it vulnerable for transposable element movement. We identified a transposable element retrotransposition into a transporter of glucosinolate defense compounds. As a consequence of this transposable element movement, the plants showed tissue-specific changes in glucosinolate profiles and levels accompanied by rewiring of glucosinolate- and defense-related transcriptional changes. As this single transposable element had strong impact on the plants’ resistance to insect herbivory, our findings highlight the potential for transposable elements to play a role in plant adaptation.

## Introduction

In the mid-20^th^ century, Barbara McClintock made the revolutionary discovery of mobile DNA, proving that genomic sequences are not necessarily stationary, but can relocate (McClintock, 1950). Such transposable elements (TEs) act either by a cut-and-paste mechanism (DNA transposons) or copying via an RNA-intermediate (retrotransposons) and they provide extensive genetic variation across kingdoms (Ross et al., 2021; Stuart et al., 2016; Wells and Feschotte, 2020; Wicker et al., 2007). Silencing of TEs is controlled by epigenetic factors, including DNA methylation (Fan et al., 2022; Matzke and Mosher, 2014; Slotkin and Martienssen, 2007; Tsukahara et al., 2009). TEs were long thought to move randomly, but recent studies have shown that insertions might not merely be a provider of random genetic variation. Recently, TEs have been proposed to play a role in adaptation based on their potential for rapid mutagenesis (Lu et al., 2021; Michael Thieme et al., 2022; Roquis et al., 2021). Changes in environmental conditions can induce global changes in DNA methylation patterns that act in controlling TE movement (Dowen et al., 2012; Van Dooren et al., 2020; Wojtyla et al., 2020). The Copia TE superfamily has shown to over-accumulate in genes which function in gene regulation making them potential players in adaptation (Lisch, 2013; Negi et al., 2016; Quadrana et al., 2019, 2016; Slotkin and Martienssen, 2007).

Studying effects of DNA methylation on trait variation is inherently challenged by the confounding effect of associated DNA sequence variation (Johannes et al., 2009, 2008). To overcome this issue, an epigenetic Quantitative Trait Loci (epiQTL) mapping population has been generated using a *decrease in DNA methylation I* (*ddm1)-*mutant in *Arabidopsis thaliana*, which is characterized by demethylation at 70% of all methylated cytosines and low levels of DNA methylations due to the mutant’s deficiency in chromatin remodeling (Vongs et al., 1993). The *ddm1* mutant allele was subsequently removed by backcrossing to Col-0 wild type (WT) and seeds from a single plant were propagated for several generations to produce lines with variation in DNA methylation patterns (Johannes et al., 2009). Part of the methylome in these epigenetic Recombinant Inbred Lines (epiRILs) are 126 stable Differential Methylated Regions (DMRs) that allow for mapping epiQTL (Colomé-Tatché et al., 2012; Cortijo et al., 2014). These epiRILs have been widely used to assess the impact of epigenetic variation on the plasticity of a vast number of traits covering plant morphology, development and defense chemistry (Aller et al., 2018; Cortijo et al., 2014; Furci et al., 2019; Johannes et al., 2009; Kooke et al., 2015; Latzel et al., 2012; Roux et al., 2011; Zhang et al., 2013). As TE movements in epiRILs are well known to occur and can be accounted for, the same epiRIL population made it feasible to compare the relative impact of TE movement and DNA methylation variation on a given trait (Cortijo et al., 2014; Johannes et al., 2009; Quadrana et al., 2019).

In a previously conducted epiQTL mapping using the *ddm1*-derived population, we identified epiRIL573 as strong outlier for the leaf accumulation of glucosinolate (GLS) defense compounds and removed it from the data set prior to further analysis (Aller et al., 2018). In this study, we focus on the outlier and localize a retrotransposition of the Copia type LTR transposon family *ATCOPIA13* to the gene encoding GLS transporter 2 (*GTR2*, ATNPF2.11, *At5g62680*). We assess the phenotypic and transcriptomic consequences of this retrotransposition event. In-depth characterization of epiRIL573 revealed regulatory networks involved in feedback regulation of GLS in roots and highlights the potential of TE retrotransposition as a mechanism enabling rapid adaptation.

## Results

### A recessive causal allele is associated with high LC GLS levels in epiRIL573

In a previous study (Aller et al., 2018), we analyzed the phenotypic response of GLS in relation to DNA methylations using a *ddm1*-derived epiRIL population (Johannes et al., 2009). GLS are defense-related compounds almost exclusively found in the order Brassicales and have been extensively studied as a model system for specialized metabolites (Andersen et al., 2013; Jensen et al., 2014; Jeschke et al., 2019; Kliebenstein et al., 2001; Nintemann et al., 2018). *A. thaliana* contains up to 40 different GLS that are classified based on their chemical structure. Indole GLS are tryptophan-derived, whereas aliphatic GLS are methionine-derived and have side chains consisting of three to eight carbon atoms. Aliphatic GLS can be further subdivided into short chain aliphatic GLS (SC GLS) and long chain aliphatic GLS (LC GLS) dependent on the length of side chain (Jensen et al., 2014).

EpiRIL573 was identified as an extreme outlier for LC GLS. The line accumulated >9 SD higher levels in rosette leaves compared to the epiRIL population mean (Figure 1). It seemed unlikely that this phenotype arose from stable methylation patterns alone, as the individual DMRs in epiRIL573 are not unique to this line, but no other epiRIL showed a similarly pronounced GLS phenotype. Instead, we hypothesized a genetic cause for the markedly higher accumulation of LC GLS.

**Figure 1:**
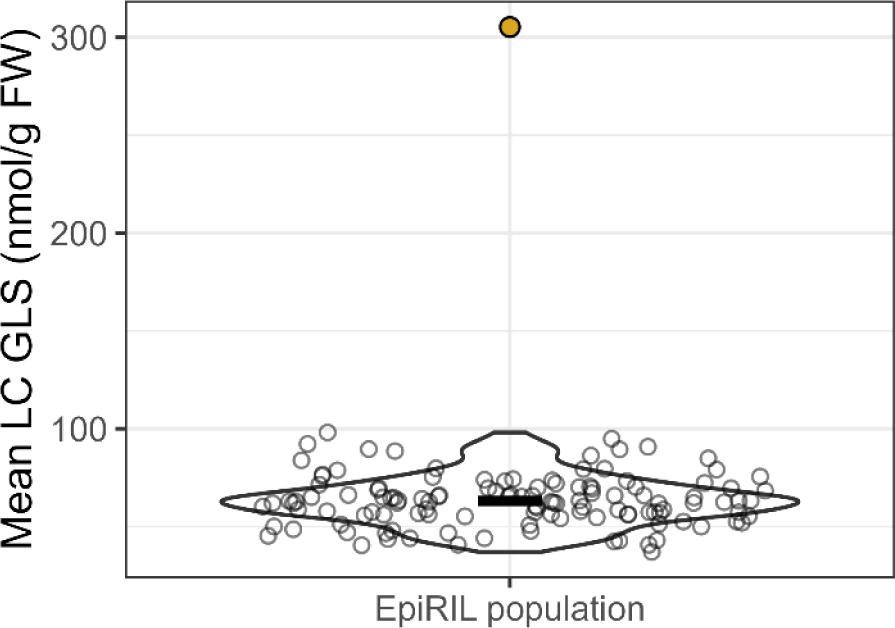
LC GLS levels in epiRIL573 compared to rest of population. Violin plot showing distribution of mean LC GLS levels/ epiRIL. Transparent points show the individual epiRIL mean level and the black line mark the population mean (excluding epiRIL573). EpiRIL573 is colored yellow. 122 epiRILs and four Col-0 WT lines were measured. Analysis was done on rosette tissue of 22-23-day old plants. n_epiRIL/WT_ = 12-18.

To address the possibility that one or more genetic alterations in epiRIL573 caused the GLS phenotype, we first tested for Mendelian segregation. We backcrossed epiRIL573 to Col-0 WT (epiRIL602) and analyzed LC GLS levels in rosette leaves of the F1 and F2 generations. F1 plants derived from reciprocal crosses accumulated LC GLS to the same levels as the parental Col-0 WT (epiRIL602), which excluded a dominant mutation as explanation for the epiRIL573 phenotype (Supplemental Figure S1A) (Aller et al., 2018). Of 259 F2 plants analyzed, 19,7% segregated with the epiRIL573 phenotype (Supplemental Figure S1C) supporting segregation of one major-effect recessive allele and potentially one or more minor-effect loci in epiRIL573 correlating with its GLS phenotype (chi-square test: X^2^= 3,9, df= 1, p-value= 0.05). No apparent morphological rosette traits co-segregated with the LC GLS phenotype (Supplemental Figure S1D).

### A retrotransposition into a GLS transporter correlates with the epiRIL573 phenotype

To identify the major-effect mutation underlying the epiRIL573 GLS phenotype, we performed bulk genome sequencing on 16 of the segregating F2 plants, eleven of which had the epiRIL573 LC GLS phenotype and five the WT phenotype (Supplemental Figure S2A). Surveying known GLS loci for genetic alterations revealed an insertion of the Copia type LTR transposon family *ATCOPIA13* (*AT2G13940*) in after the second exon of *GLS transporter 2* (*GTR2*, *AT5G62680*) (Nour-Eldin et al., 2012) (Supplemental Figure S2B).

We further validated the *GTR2* TE-insertion by genotyping a region spanning the TE insertion in epiRIL573 and compared to a T-DNA mutant *gtr2-1* line (Figure 2B) (Nour-Eldin et al., 2012). All tested epiRIL573 plants carried the TE insertion, which was absent from Col-0 WT *GTR2* as well as in *gtr2-1* (Figure 2B). We additionally genotyped F2 segregants which showed that the LC GLS phenotype co-segregated with the TE insertion (Supplemental Figure S3).

**Figure 2:**
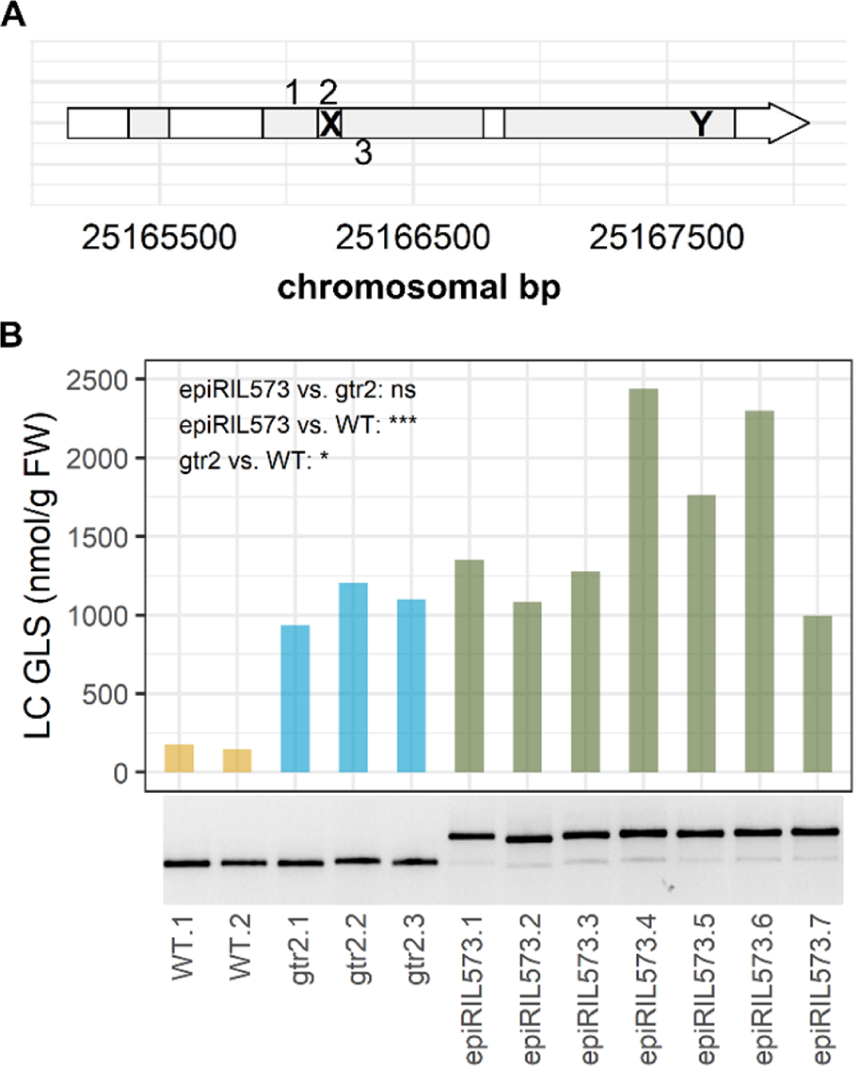
Amplification of TE insertion. A) Model showing the *GTR2* gene *(At5g62680)* and its position on chromosome 5 (reverse strand), exons are colored grey. “X” represents site of TE (*At2g13940*) insertion in epiRIL573. “Y” represents site of T-DNA insertion in *gtr2-1* SAIL mutant line. Numbers show primer annealing sites. B) Plot shows LC GLS levels in the three genotypes. Significant differences were assessed using a games-Howell adjusted post hoc test on the model LC ∼Genotype. The gel in the lower panel shows the amplicons obtained using primer combination 1+2 (amplification of ∼690 bp in presence of the TE insertion) and primer combination 1+3 (amplification of 467 bp in the absence of TE insertion). PCR samples were pooled before gel electrophoresis.

TE mobility is activated upon mutation in the *DDM1* gene with continued activation after restoration of WT *DDM1* (Kato et al., 2004). As the epiRIL population was generated using the *ddm1-2* mutant, epiRIL573 potentially carries additional genetic rearrangements. To identify other potential retrotranspositions, we performed a global TE search on epiRIL573 using publically available epiRIL genome sequencing data (Supplemental Figure S4A) (Quadrana et al., 2019). We filtered for homozygous TEs in epiRIL573 and identified one additional TE movement besides the retrotransposition into *GTR2*. A TE from the DNA transposon VANDAL21 family, *AT2TE42810* has moved into *ATSYTF* (*AT3G18370)* in epiRIL573. *AT3G18370* encodes a C2 domain-containing protein putatively associated with leaf formation (Huercano et al., 2022). To test whether this TE insertion was connected to the LC GLS phenotype, we did an additional global TE search in our segregating F2 plants grouped by their phenotypic segregation (Supplemental Figure S4B). We could not detect the insertion into *ATSYTF* in plants with WT phenotype. In F2 plants with the epiRIL573 phenotype, we did detect the insertion. However, the allele frequency was 0,63 and the insertion did not co-segregate with the LC GLS phenotype. This points to the insertion into *ATSYTF* having no apparent impact on GLS accumulation in epiRIL573.

To test for phenotypic rescue upon reintroducing WT *GTR2*, we complemented epiRIL573 and *gtr2-1* with WT *GTR2* (Supplemental Figure S5). It was recently found that successful complementation partly relies on the length of the putative promoter included (Sanden et al., Accepted for publication). Here, we used a ∼2kb native promoter and a ∼8kb native promoter driving expression of GTR2 CDS and gene, respectively. T1 transformants were confirmed by genotyping and analyzed for GLS. For both constructs, epiRIL573 showed a partly rescued phenotype upon complementation, which was more pronounced using the 8kb promoter (Supplemental Figure S5B). *gtr2-1* also showed a partial rescue, but not to the same degree as for epiRIL573. The partial complementation suggests that neither construct comprised all regulatory sequence elements to fully restore *GTR2* expression levels. Nevertheless, as this is the case for both epiRIL573 and *gtr2-1*, the observed complementation supported that the GLS phenotype of epiRIL573 is associated with the disruption of *GTR2*.

### EpiRIL573 phenotypically mimics *gtr2-1*

GTR2 is a plasma membrane-localized H+/GLS symporter, which mediates the uptake of GLS from the apoplast (Madsen et al., 2014; Nour-Eldin et al., 2012). Retrotransposition of *ATCOPIA13* into the second intron of the *GTR2* gene (Figure 2A) likely renders the protein completely non-functional, as it interrupts the GTR2 protein sequence after 127 of 617 amino acids. In the previously described *gtr2-1* mutant, insertion of a T-DNA into the last exon abolishes only 41 amino acids at the C-terminus (Nour-Eldin et al., 2012, Supplemental Figure 4). We therefore sought to further phenotypically compare the newly identified *gtr2* mutant with *gtr2-1*.

Upon analysis of leaf GLS, epiRIL573 and *gtr2-1* both displayed markedly higher LC GLS levels in rosette tissue, as observed for *gtr2-1* before (Figure 2A, Figure 3A) (Hunziker et al., 2021). The mutant lines accumulated SC GLS to WT levels (Figure 3B). EpiRIL573 and *gtr2-1* further showed a similar trend towards higher levels of indole GLS in leaves (Figure 3C). In seeds, epiRIL573, but not *gtr2-1* had significantly lower levels of SC GLS than WT (Figure 3D). Both mutant lines had significantly lower levels of LC GLS and indole GLS in seeds compared to WT (Figure 3 E-F). Our analysis confirmed the previously published data on *gtr2-1*, which suggested an increase in total aliphatic and indole GLS in rosette tissue and a reduction the same GLS in seeds (Nour-Eldin et al., 2012). We thus concluded that epiRIL573 displays the same leaf and seed GLS phenotypes as *gtr2-1*.

**Figure 3:**
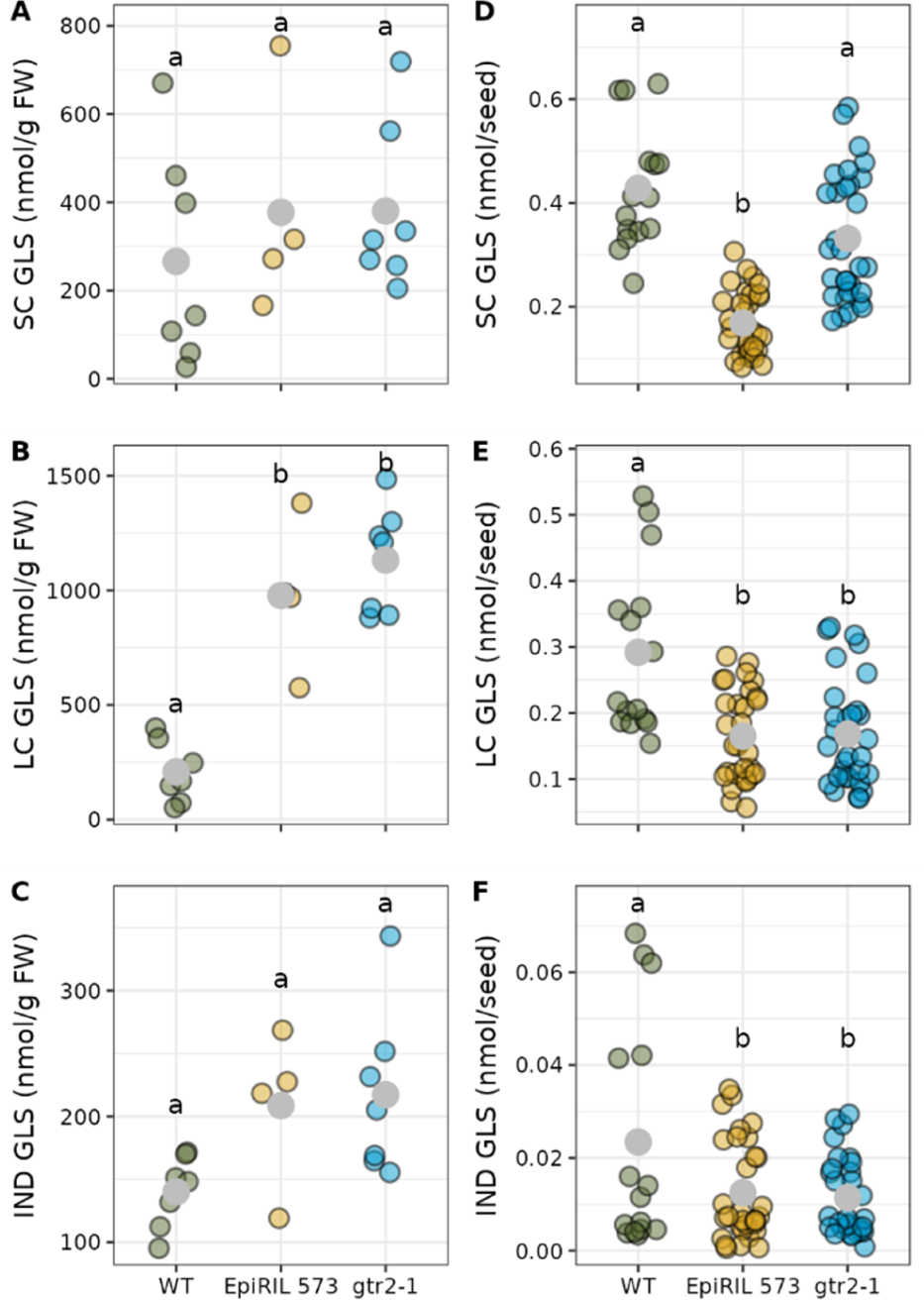
GLS levels in epiRIL 573 vs. *gtr2-1* in rosette tissue and seeds. Grey points mark mean levels. Data was analyzed with a post hoc Tukey test on model GLS∼ Genotype+ error(Biorep) with significance threshold of pval < 0,05. A-C) Rosette tissue from 29-day-old plants, n_genotype_= 4-7. D-F) Seeds, n_genotype_= 8-16. GLS, glucosinolate; SC, short chain; LC, long chain; IND, indole; WT, pooled data from the two corresponding WTs.

### *S. littoralis* feeding pattern indicates an altered GSL distribution in rosettes of epiRIL573

The spatial distribution of GLS in *A. thaliana* is critical for the defensive function of the glucosinolate-myrosinase system. In Col-0 WT, younger rosette leaves contain higher levels of GLS than older leaves (Brown et al., 2003; Hunziker et al., 2021). Younger leaves have further been shown to possess higher myrosinase activity, i.e. potential for more rapid release of GLS activation products (Burow et al., 2007); and to release a different profile of GLS-derived bioactive compounds (Wentzell and Kliebenstein, 2008). In bioassays with *Spodoptera littoralis*, larvae fed preferentially on older rosette leaves of Col-0 WT plants. In the *gtr2-1* knockout, rosettes showed an equal distribution of GLS among leaves of different developmental stages and larvae of *S. littoralis* fed equally on young and old leaves of *gtr2-1* (Hunziker et al., 2021).

When we tested the herbivore feeding preference of *S. littoralis* on epiRIL573 in comparison to *gtr2-1* and their corresponding WTs (epiRIL602 and segregating Col0, respectively), the feeding pattern on epiRIL573 mimicked that on *gtr2-1* (Figure 4). In the absence of a functional GTR2, *S. littoralis* no longer preferred mature rosette leaves over young leaves as seen for the corresponding WT lines (Figure 4B). The weight of *S. littoralis* caterpillars at the end of the feeding period was not strongly affected by plant genotype (Figure 4A). We conclude that the larvae show similar feeding preferences on epiRIL573 and *gtr2-1*, which suggests that epiRIL573 has the same altered distribution of GLS within the rosette as *gtr2-1*.

**Figure 4:**
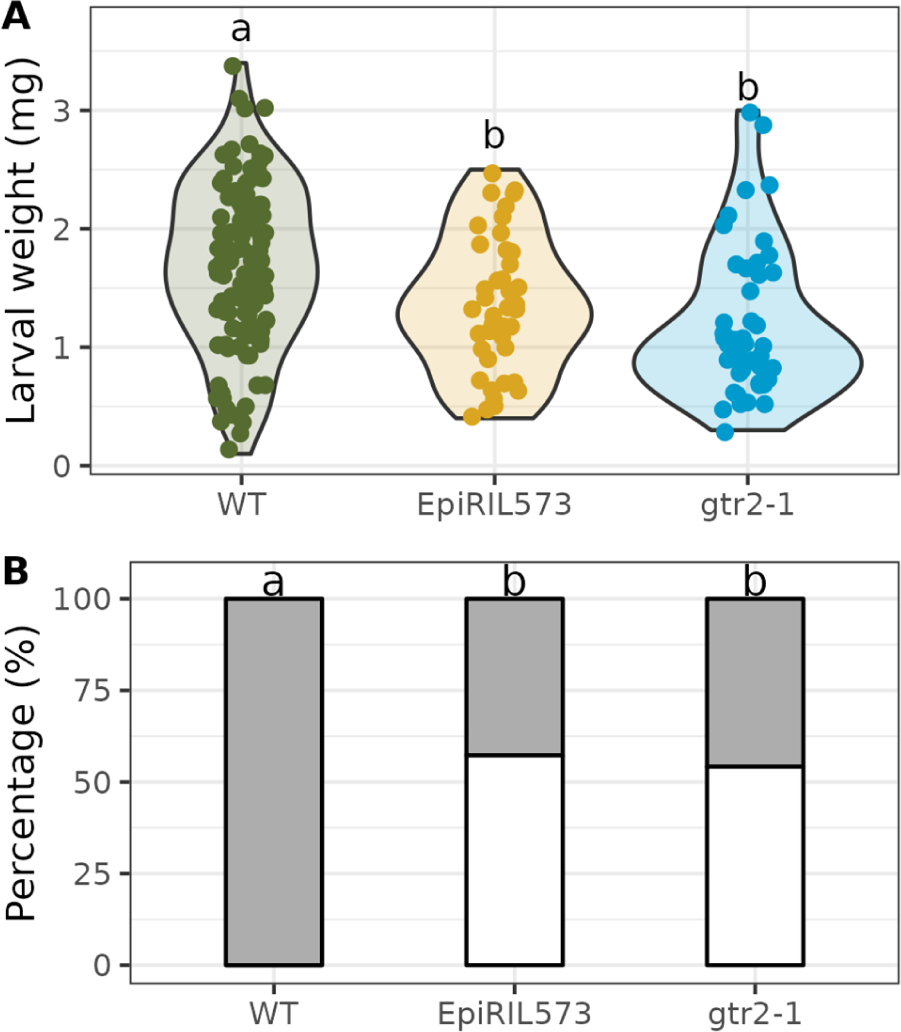
*S. littoralis* feeding assay. The assay was conducted on epiRIL573, *gtr2-1* and the corresponding Col-0 WTs. A) Weight of *S. littoralis* caterpillars after feeding on the different lines. Data was analyzed with a post hoc Tukey test on model Weight∼Genotype+ error(experimental replicate) with significance threshold of pval < 0,05. B) Fraction of feeding damage on mature leaves (grey) versus young leaves (white) within a rosette. Data was analyzed with a Fisher’s exact test and letters indicate significant differences (p-val <0.05).

### Altered GLS distribution EpiRIL573 reveals feedback regulation in roots

LC GLS are synthesized to a larger extent in roots than in rosette tissue of *A. thaliana* (Andersen et al., 2013). In roots of the *A. thaliana* double mutant lacking a functional GTR2 and its closest homologue GTR1 (ATNPF2.10), GLS are not retained in the root and travel upwards via the xylem. This increases the flux from root to rosette tissue and as a result, aliphatic GLS levels are high in rosette tissue and low in roots (Andersen et al., 2013; Hunziker et al., 2021). To assess the impact of GTR2 on the root GLS phenotype, we analyzed roots of epiRIL573 representing a *gtr2* single mutant (Figure 5A). SC and LC GLS were significantly lower in roots of epiRIL573. Indole GLS were significant changed. The almost depletion of aliphatic GLS in epiRIL573 compared to its corresponding Col-0 WT due to pronounced decrease in SC and LC GLS levels (Figure 5B).

**Figure 5:**
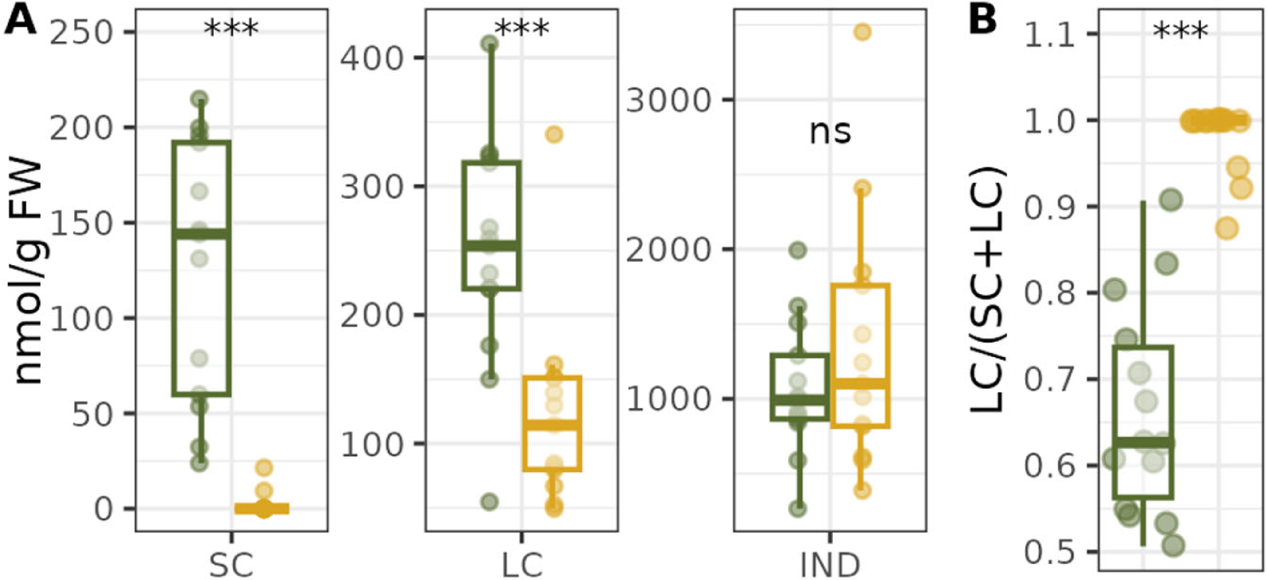
Root GLS levels profiles. Root tissue of 27-day-old epiRIL573 and WT plants. N= 14-20. WT and epiRIL573 were compared using a Bonferroni adjusted t-test. Green = WT, yellow = EpiRIL573 A) GLS levels of SC GLS, LC GLS, IND GLS. Stars denote a p-value < 0.001. B) The fraction that LC GLS constitute of total aliphatic GLS in root tissue. GLS were also measured in the corresponding rosettes (Supplemental data file S1).

Changes in GLS levels and composition have been shown to feedback regulate GLS biosynthesis and the surrounding regulatory networks in *A. thaliana* leaves and seedlings (Burow et al., 2015; Francisco et al., 2016; Jensen et al., 2015; Jeschke et al., 2019; Wentzell et al., 2007, 2007). We exploited epiRIL573, a mutant with a strong GLS phenotype and an intact regulatory and biosynthetic machinery, to identify potential regulatory responses to the increased efflux of GLS from the root. To study regulatory responses, we performed RNA sequencing on root tissue of epiRIL573 and WT plants (Supplemental Figure S6). Transcriptome sequencing revealed 888 differentially expressed genes (DEGs) in epiRIL573. Of these, 439 DEGs were upregulated and 449 DEGs were downregulated. Neither AtSYTF, which also carries a TE insertion in epiRIL573, nor genes co-expressed with AtSYTF (Obayashi et al., 2022) were among the DEGs.

To gain insight to processes impacted in epiRIL573, we enriched for GO terms in up- and downregulated genes (Table 1, full table is found in Supplemental data file S2). Upregulated genes involved in wounding, GLS biosynthesis, jasmonic acid (JA) signaling, and the regulation of defense responses were found to be overrepresented. A large proportion of the downregulated genes were associated with different catabolic processes, including genes encoding GLS catabolic enzymes.

**Table 1:** Top 5 GO Enrichments for Biological Processes in up- and down regulated DEGs in epiRIL573 vs. WT root tissue. (for full lists, see supplemental data file S2)

|  |  |  |
| --- | --- | --- |
| Up regulated DEGs |  |  |
| GO term | Count | FDR |
| ~response to wounding | 73 | 3,15E-25 |
| ~glucosinolate biosynthetic process | 25 | 6,98E-19 |
| ~response to jasmonic acid | 44 | 1,11E-14 |
| ~protein folding | 17 | 2,28E-06 |
| ~regulation of defense response | 34 | 3,38E-06 |
| Down regulated DEGs |  |  |
| ~organonitrogen compound catabolic process | 41 | 1,07E-22 |
| ~cellular catabolic process | 48 | 1,34E-20 |
| ~response to light intensity | 30 | 8,09E-14 |
| ~response to absence of light | 12 | 1,02E-07 |
| ~leucine catabolic process | 6 | 4,63E-06 |

We then integrated the genome-wide data sets on epiRIL573 by overlaying the map of stable DMRs (Colomé-Tatché et al., 2012), root DEGs, root expression of GLS-related genes specifically and sites of TE transpositions (Figure 6). The stably inherited DMRs in epiRIL573 are mainly hypermethylated (Figure 6, outer track, blue color), with the exception of one region on chromosome 3 that is hypomethylated (Figure 6, outer track, yellow color). The same region displayed a burst of upregulated genes in the root (Figure 6, middle track), which could possibly be connected with a role of cis-acting transcriptional repression by the methylations in this region that were not maintained in epiRIL573 (He et al., 2022; Zilberman et al., 2007). An additional patch of genes highly expressed in epiRIL573 was, however, found on chromosome 1. The two retrotranspositions that occurred in epiRIL573 did not coincide with regions showing high density of DEGs with high fold differences.

**Figure 6:**
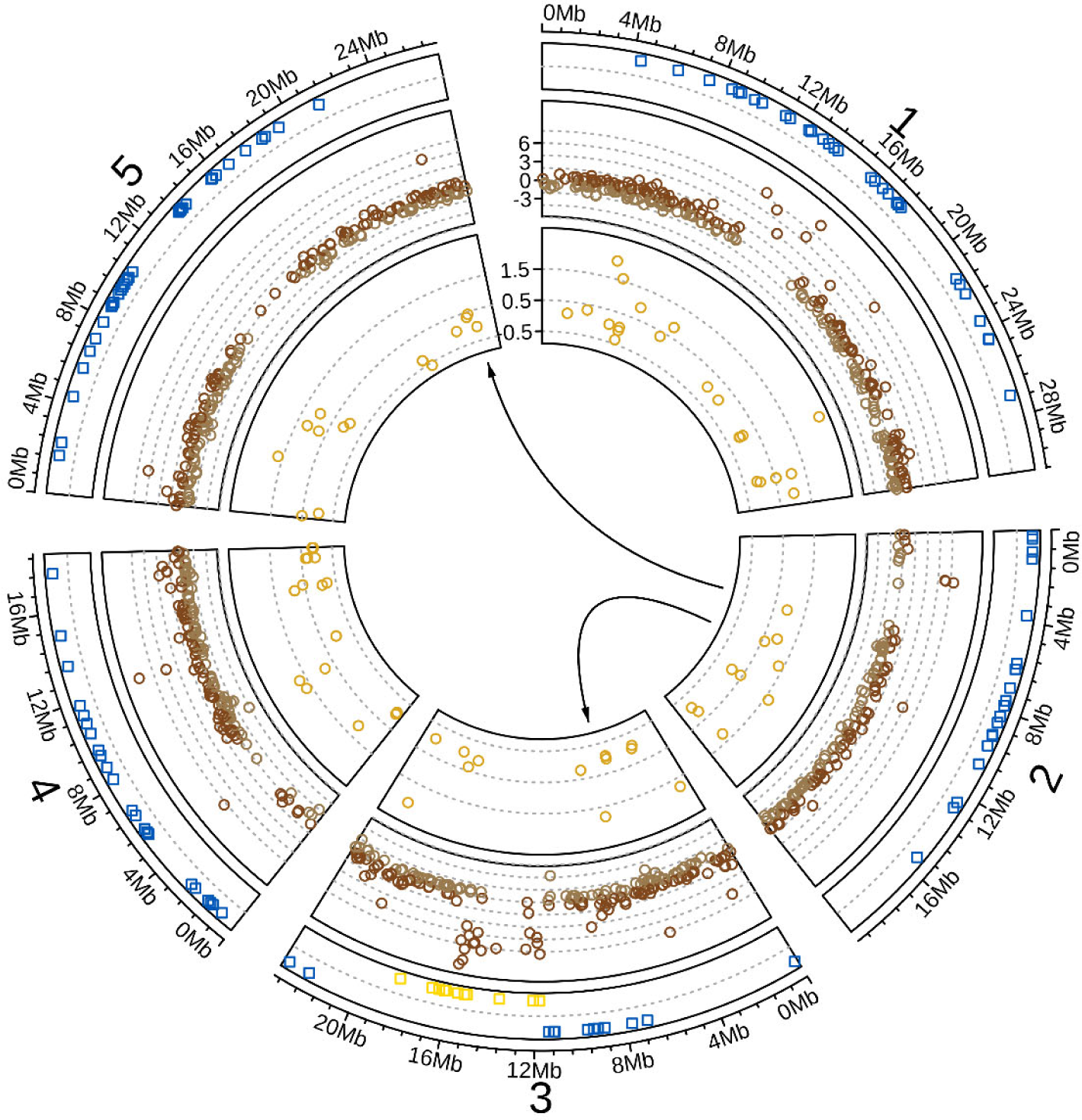
Omics integration of epiRIL573. Circular plot overlapping data as follows: Outer track shows the stably inherited DMRs in epiRIL573 with blue regions denoting hypermethylated regions and yellow regions denoting hypomethylated regions. Middle track shows log2FC of significantly DEGs in root (padj < 0.05). Dark brown marks upregulated genes and light brown marks downregulated genes. Inner track shows log2FC expression of GLS pathway genes irrespective of significance. Arrows show TE transpositions. The arrow from chromosome 2 to 5 marks the retrotransposition into *GTR2 (AT5G62680)*. The arrow from chromosome 2 to chromosome 3 marks the movement into *ATSYTF (AT3G18370)*.

Genes involved in regulation, biosynthesis, catabolism, and transport of GLS are plotted as their log2FC in epiRIL573 irrespective of significance (Figure 6, inner track). Whereas biosynthetic genes for some specialized metabolites are arranged in gene clusters (Polturak and Osbourn, 2021), GLS-related genes are scattered across the *A. thaliana* genome (Kliebenstein, 2009). Changes in GLS gene expression were also not location specific. We zoomed further in on GLS-related genes and grouped them by their associations to different GLS processes (Figure 7, Supplemental data file S3). Despite the approximately same number of genes globally being up and down regulated, GLS-genes showed mostly upregulation. The biggest transcriptional differences were seen for regulators and enzymes involved in the biosynthesis of aliphatic GLS (Figure 7). Some genes related to indole GLS were also differentially expressed in roots of epiRIL573, although the corresponding metabolites accumulated to WT levels in the root (Figure 5C). Further, transcript levels of the myrosinase *TGG2* (*AT5G25980*) and the Nitrile Specifier Protein *NSP5* (*AT5G48180*) differed significantly between epiRIL573 and the WT, suggesting fine-tuning of GLS catabolism in the absence of a functional GTR2. Despite the TE insertion, GTR2 was expressed at low levels, which could be attributed to the two first exons of the gene still being transcribed (Supplemental Data S3, Supplemental Figure S7).

**Figure 7:**
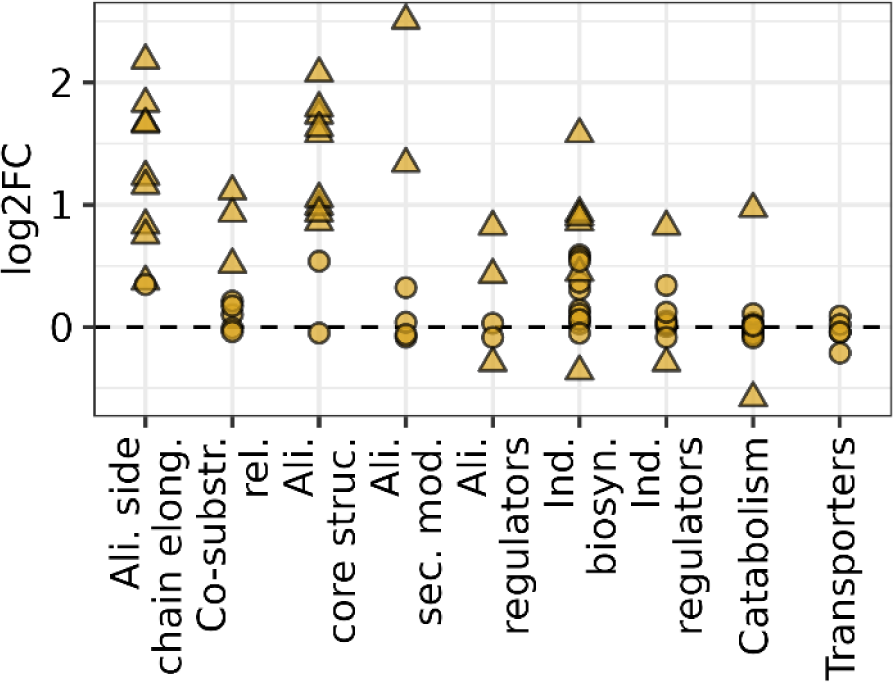
Gene expression of GLS-related genes in epiRIL573 root tissue. Log2FC of gene expression in epiRIL573 vs. WT is grouped by the GLS process genes are associated with (for individual gene expression data, see Suppl. Data File S2). Genes that were significantly altered in epiRIL573 vs. WT are marked as a triangle and insignificantly altered genes are marked as a circle (padj < 0.05). Ali. Side chain elong.= Aliphatic side chain elongation, Co-substr. rel= Co-substrate related, Ali. Core struc.= Aliphatic core structure, Ali. Sec. mod = Aliphatic secondary modifications, Ali. regulators= Aliphatic regulators, Ind. biosyn= Indolic biosynthesis, Ind. regulators= Indolic regulators.

A large proportion of GLS-related genes was differentially expressed in roots of epiRIL573 as also reflected by the GO term enrichment analysis (Table 1), which was most pronounced for genes controlling the accumulation of aliphatic GLS (Figure 7).

## Discussion

Assessing the impact of epigenetic regulation on quantitative variation in a given trait has proven difficult because of the confounding genetic variation that affects the same trait. The *ddm1*-derived epiRIL population was developed as an approach to overcome this issue (Johannes et al., 2009). This population was designed to have a common genetic background with significant variation in DNA methylation patterns, allowing for trait variation to be attributed to DMRs. However, TE movement is a well-known consequence of manipulating DNA methylation, and transpositioned TEs have previously been identified in epiRILs with movements tracked back to the early steps in generation of the population (Cortijo et al., 2014). These early TE movements did not compromise the correlation between phenotypes and DNA methylations to a large extent as the impact on trait variation could largely be attributed to the associated epiQTL and only to a lesser degree to the identified TE movements (Cortijo et al., 2014). Genome sequencing of 107 epiRILs from the *ddm1*-derived epiRIL population more recently indicated that many additional TE transpositions occurred later during propagation of the epiRILs and that TE movement can still occur eight generations after reconstituting *DDM1* (Quadrana et al., 2019; Tsukahara et al., 2009).

Our data on epiRIL573 illustrates an example of a single TE retrotransposition in an individual epiRIL, which had a major impact on the investigated trait. EpiRIL573 accumulated approximately nine times higher LC GLS levels in rosette tissue compared to the mean accumulation in the epiRIL population (Fig. 1) (Aller et al., 2018). As we otherwise found the impact of variation of DNA methylations on GLS trait variation to be relatively low, we tested the hypothesis of a genomic rather than purely epigenomic alteration. We genome-sequenced plants from the F2 generation after backcrossing to WT and identified two TE transpositions in epiRIL573, but only the insertion of an LTR retrotransposon (*At2G13940*) into the GLS transporter gene *GTR2 (At5G62680)* co-segregated with the high LC GLS phenotype. Given the strong phenotypes observed in plants homozygous for the TE insertion in *GTR2*, this retrotransposition event is very unlikely to have happened in other epiRILs, but represents single seed descent of the lines during the generation of the population (Johannes et al., 2009; Quadrana et al., 2019).

The phenotypic impact of the TE transposition into *GTR2* in one epiRIL was substantial enough to affect DMR mapping. Only after removing epiRIL573 from the data set, a DMR that significantly correlated with variation in LC GLS levels in leaves could be mapped (Aller et al., 2018). Among the 107 epiRILs that were recently genome-sequenced, 95% displayed at least one and up to 97 TE transpositions that were not carried through from the *ddm1* parent (Quadrana et al., 2019). We can therefore not rule out that additional TE movements in other epiRILs in the population contribute smaller quantitative shifts in GLS trait variation and thereby had an impact on the mapped DMRs. Minor effect mutations induced by TE movement would not have rendered the respective epiRILs as extreme outliers and would thus not have been removed for analysis.

GTR2 and its close homologues GTR1 and GTR3 (ATNPF2.9) mediate uptake of GLS across the plasma-membrane and thereby impact long-distance transport of GLS. Before the onset of flowering, GTRs contribute to controlling the root-to-shoot distribution of aliphatic and indole GLS (Andersen et al., 2013; Jørgensen et al., 2017). Plants lacking both GTR1 and GTR2 have previously been reported to accumulate high levels of all GLS, but especially LC GLS in leaves (Andersen et al., 2013; Jørgensen et al., 2017; Madsen et al., 2014). The *gtr2-1* single mutant is a major factor for over-accumulation of LC GLS in leaves irrespective of their developmental stage (Hunziker et al., 2021). epiRIL573, which had been identified based on its high GLS levels in leaves, phenotypically mimicked *gtr2-1* (Figure 3), supporting the conclusion that all GLS phenotypes observed in epiRIL573 under the conditions tested can be attributed to the retrotransposition event that disrupted *GTR2*. Reintroducing the GTR2 coding sequence under the control of ca. 2kb of its putative promoter did not fully restore WT levels of LC GLS Complementation with the construct including 8kb promoter and the genomic GTR2 sequence largely rescued the GLS phenotype (Supplemental Figure S5), suggesting that regulatory elements in the distal promoter and/or introns are critical for *GTR2* regulation at the transcriptional level.

Double mutants in *A. thaliana* and *Camelina sativa*, devoid of GTR1 and GTR2, accumulate only very low levels of aliphatic GLS (Andersen et al., 2013; Hölzl et al., 2023), whereas levels of indole GLS were found be higher than in WT roots (Andersen et al., 2013; Jørgensen et al., 2017). In roots of epiRIL573, representing a *gtr2* single mutant, we observed substantially lower levels of aliphatic GLS compared to WT (Figure 5). Although GTR2 shows no preference for aliphatic GLS when expressed in *Xenopus laevis* oocytes (Jørgensen et al., 2017), indole GLS levels were unaffected. This could either be due to spatial differences in biosynthesis and storage between aliphatic and indole (Nintemann et al., 2018) GLS and/or be explained by different mechanisms of feed-back regulation.

To gain insight into the transcriptional response to the TE retrotransposition into *GTR2* and the resulting root GLS phenotype, we sequenced RNA from root tissue of epiRIL573 and its corresponding WT. Known GLS related genes showed a distinct pattern (Figure 7). Almost all genes encoding enzymes involved in the chain elongation of methionine and the subsequent core structure formation of aliphatic GLS were significantly upregulated in epiRIL573. Likewise, genes critical for the supply of 3’-phosphoadenosine 5’-phosphosulfate (PAPS) as sulfur donor for methionine chain elongation (Møldrup et al., 2011; Yatusevich et al., 2010) showed elevated expression levels in epiRIL573. In contrast, biosynthetic genes specific to indole GLS, genes involved in GLS catabolism and GLS transporters were largely unchanged in expression levels. Thus, the increased efflux of aliphatic GLS from the roots to the shoots is accompanied by transcriptional feedback activation that is specific to aliphatic GLS biosynthesis.

The elevated expression levels of genes involved in aliphatic GLS biosynthesis were highly consistent. The log2FC in epiRIL573 compared to WT varied around an average of 1.3 and did not exceed 2.5 (Supplemental data file S3). These quantitatively minor differences correspond well with previous studies on transgenic lines expressing over-expressed copies of GLS regulators, where moderate increases in transcript levels of GLS pathway genes led to significant higher GLS accumulation (Li et al., 2013; Sønderby et al., 2007). The upregulation of aliphatic GLS biosynthesis in epiRIL573 can thus be considered a strong GLS response.

Among the upregulated genes in the root transcriptome of epiRIL573, several other defense-related GO terms that were also found to be overrepresented, among them JA signaling (Table 1). Upregulated genes involved in JA biosynthesis and signaling included *MYC2* (*AT1G32640*), *JAZ2* (*AT1G74950*), *JAZ5* (*AT1G17380*), *JAZ6* (*AT1G72450*), *LOX3* (*AT1G17420*), AOS (*AT5G42650*), and OPCL1 (*AT1G20510*).

JA signaling has a large effect on the regulation of both aliphatic and indole GLS (Dombrecht et al., 2007; Hirai et al., 2007; Mewis et al., 2005). The R2R3 MYB transcription factors that act as positive regulators of GLS biosynthesis are functionally dependent on interaction with a MYC bHLH transcription factor (Frerigmann et al., 2014; Millard et al., 2019), and consequently, triple mutants lacking MYC2, MYC3, and MYC4 are almost completely devoid of GLS (Schweizer et al., 2013). In rosette leaves, the GLS regulatory network has further been shown to feedback regulate JA biosynthesis and signaling (Burow et al., 2015). The upregulation of genes of the JA signaling network suggests that elevated JA signaling is needed to channel resources into GLS biosynthesis and thereby increase GLS production. Altered resource allocation in epiRIL573 is further reflected by the down-regulation of leucine biosynthesis genes, which share the isopropylmalate dehydrogenase *IPDMH1* with the methionine chain elongation pathway in GLS biosynthesis (He et al., 2009). Alternatively, roots of epiRIL573 might perceive the low levels of GLS similar to a biotic interaction that triggers GLS activation and induces JA signaling. Taken together, TE-induced disruption of *GTR2* in epiRIL573 did not only have a major impact on GLS accumulation, but also strongly affected aliphatic GLS at the transcriptional level by rewiring the surrounding regulatory networks.

## Conclusions

Uncoupling epigenetic and genetic mechanisms to evaluate their individual and joint impact on trait variation has proven challenging. The *ddm1*-derived epiQTL mapping population allows correlation of stably inherited DMRs with trait variation. These correlations must, however, be interpreted with caution due to the increased rate of TE transposition in the population. epiRIL573 exemplifies that a single TE movement during the single seed descent can cause major defense-related phenotypes. The in-depth characterization of this line illustrates the potential of TE transposition for rapid adaptation to environmental changes. By encompassing both a role in creating natural variance and taking part in active adaptation, TEs could in theory bridge Darwin’s evolutionary model to the thoughts on active acquisition proposed by Lamarck.

## Materials and Methods

### Germplasm

122 *A. thaliana* epiRILs generated in the Col-0 background and four Col-0 WT lines (Colomé-Tatché et al., 2012; Johannes et al., 2009) were purchased from the Versailles Arabidopsis Stock Center, Institut Jean Pierre Bourgin (INRAE, n.d.; Johannes et al., 2009). EpiRIL “35RV573” is referred to as epiRIL573. The corresponding Col-0 WT “35RV002” was used as control and is here referred to as epiRIL602. The *gtr2-1* T-DNA insertion line, SAIL_20_B07, was previously described (Nour-Eldin et al., 2012). Data from epiRIL602 and the Col-0 WT grown-along with gtr2-1 were pooled after statistically ensuring their similarity.

### Plant Handling and Growth Conditions

Seeds were vapor-sterilized for 2 h by exposure to a mixture of 18 mL bleach (Sodium hypochlorite 14%) and 2 mL HCl (37%). Seeds were sown on a potting soil (Pindstrup nr. 2, Pindstrup Mosebrug A/S, Denmark) and sand mixture (3:1), cold-stratified for 4-6 days at 4°C in the dark and subsequently grown in a light chamber set to 80-120 μE/ (m^2^*s), 16 h light, 21°C, and 70% relative humidity for 21-22 days. For GLS analysis, plant tissue was harvested 21-29 days after stratification as indicated in figure legends. Root tissue was obtained by dipping in miliQ water. Roots and rosettes were weighed separately. For RNA-sequencing, rosette tissue was harvested from 23-day old plants and root tissue from 27-day old plants. The plant material was snap-frozen in liquid nitrogen and stored at −80°C.

### S. littoralis feeding assay

For insect bioassays, plants were grown under short-day conditions (10h light, 150 μE/ (m2*s), 21°C, 70% relative humidity) for ca. 4 weeks. The assays were carried out as previously described (Hunziker et al., 2021).

### GLS Analysis

GLS were extracted and analyzed as desulfo-GLS (dsGLS) as described previously (Crocoll et al., 2016; Kliebenstein et al., 2001). Sigma-Aldrich Millipore 96 well filter plates, cat.no. MSHVN45 were charged with 45 mg DEAE Sephadex A25 and 300 ml of water per well and equilibrated at room temperature (RT) for minimum 2 h. The water was removed using a vacuum manifold (Millipore). Tissue was harvested into with 300 μl 85% (v/v) methanol (HPLC grade) containing 5 μM *p*-hydroxybenzyl GLS (pOHb; PhytoLab, cat. No. 89793) as internal standard and homogenized with two stainless steel balls (diameter 3.5 mm) by shaking for 2 min at a frequency of 30 Hz in a Mixer Mill 303 (Retsch, Haan, Germany). Single seeds were analyzed using 1 nmol internal standard. Samples were centrifuged and the supernatants were applied to the filter plates followed by vacuum filtration for 2-4 s. Sephadex material was washed with 2x 100 µl 70% methanol (v/v) and 2x 100 µl water and briefly centrifuged before addition of 20 µl of sulfatase solution per sample (Crocoll et al., 2016). After incubation at (RT) overnight, dsGLS were eluted with 100 µl water. dsGLS were analyzed by LC-MS/MS on an Advance UHPLC system (Bruker, Bremen, Germany) equipped with a Kinetex® XB-C18 column (100 × 2.1 mm, 1.7 μm, 100 Å, Phenomenex, USA) coupled to an EVOQ Elite TripleQuad mass spectrometer (Bruker, Bremen, Germany) equipped with an electrospray ionization source (ESI) as previously described (Crocoll et al., 2016). Glucosinolates were quantified relative pOHb using experimentally determined response factors with commercially available standards in a representative plant matrix.

### Segregation Analysis

Reciprocal crosses were carried out with epiRIL573 and Col-0 WT (epiRIL 602). Rosette tissue from 80 F1 plants was analyzed for GLS together with four plants of grown-along epiRIL573 and WT controls. We propagated eight plants from the F1 generation (four plants being maternally derived from WT and four from epiRIL573) representing the higher and lower spectrum of LC GLS distribution. Their progeny together with the progeny of two plants from each parental line grown along with the F1 plants were analyzed for segregation over four experimental rounds. In each round, we analyzed 2-6 biological replicates of control plants and 5-12 replicates of F1 plants derived from crossing. A total of 259 F2 segregants were analyzed for GLS.

### DNA library preparation and sequencing

Rosette and root tissue was harvested from 16 plants from the F2 generation (maternally derived from WT), five plants with a WT LC GLS phenotype and 11 displaying the epiRIL573 LC GLS phenotype. The frozen tissue was ground 5x 30 seconds at 30 Hz in a Mixer Mill 303 (Retsch, Haan, Germany). 1 mL 37°C sterile extraction buffer (50 mM Tris pH 8, 200 mM NaCl, 2 mM EDTA, 0,5% (w/v) SDS) was added to each sample and the samples were incubated at 37°C for 20 min with occasional inversion. 500 μL chloroform/phenol/isoamyl alcohol (Sigma P3803) were added, samples were mixed and then centrifuged at 6000 rpm for 10 min. The aqueous phase was transferred to a new tube and 500 μL chloroform/isoamyl alcohol (Sigma C0549) were added. Samples were mixed and centrifuged at 6000 rpm for 10 minutes. The aqueous phase was moved to new tubes and 600 μL isopropanol were added for precipitation at 20°C overnight. DNA was pelleted by centrifugation at 6000 rpm for 10 min, the supernatant was removed, and the pellet was rinsed twice with cold 70% ethanol and subsequently dried at RT DNA pellets were re-solubilized in 50 μL ddH_2_0 at RT, and then precipitated with 1 mL 96% ethanol by centrifugation at 2000 rpm for 10 min. The supernatant was removed, the pellet dried at RT and finally re-solubilized in 30 μL ddH20 at RT. DNA quality was assessed using a ThermoFisher Nanodrop and DNA concentration using ThermoFischer Qubit Fluorometric Quantification with the Qubit dsDNA HS Assay Kit (Thermo Fischer, Catalogue number: Q32851).

50 µL DNA (10 µg/ml) was sheared using a sonicator (QSonica, Q800R3) with amplitude setting of 25%, a pulse frequency of 10 s on and 10 s off for 3,5 min at 3°C. Sheared DNA size of approximately 500 bp was validated by by gel-electrophoresis.

DNA libraries were made with an Illumina Neoprep system using the TruSeq Nano DNA Library Prep Kit (NP-101-1001). Sequencing was performed with an Illumina MiSeq system and the MiSeq Reagent Kit v3 (MS-102-3003). The quality of the output bam files was assessed with the program FastQC (“FastQC,” 2015) and the data was further analyzed with Integrative Genomics Viewer (IGV) (Thorvaldsdottir et al., 2013).

### Genotyping

DNA for genotyping was extracted with following CTAB-based method (Clarke, 2009). Rosette tissue was harvested, snap-frozen in liquid nitrogen and ground 2x 30 seconds at 30 Hz in a Mixer Mill 303 (Retsch, Haan, Germany) followed by 1 min centrifugation at 3000 rpm. 200 μL CTAB buffer (2% (w/v) cetyl-trimetyl-ammonium bromide, 1,4 M NaCl, 100 mM Tris HCl pH 8, 20 mM EDTA) was then added per sample, samples were vortexed and incubated at 65°C for 1 h. After cooling, 200 μL chloroform were added and samples were vortexed and spun at 2000 rpm for 15 min. 125 uL of the aqueous phase were as transferred to 125 uL isopropanol and mixed. Samples were then centrifuged for 15 min at 6000 rpm. Pellets were washed with 150 μL EtOH and spun for 5 min at 6000 rpm. Supernatants were removed and the dried pellets were then resuspended in 100 μL milliQ water.

For genotyping of epiRIL573 and WT plants, we amplified actin using following primers: Actin-F, 5’-ACATTGTGCTCAGTGGTGGA-3’ and Actin-R, 5’-TCATACTCGGCCTTGGAGAT-3’ leading to an amplification product of 288 bp. Primer sequences to amplify a region spanning the TE retrotransposition in GTR2 were as follows: A-FP, 5’-TAGACCGATCGTCCAACTGAC-3’, A-R1, 5’-GGTCGATACAAGACTCTAAGTGTC-3’. These primers only amplified the GTR2 region in the absence of a TE insertion as the insertion size compromised the PCR. In absence of insertion, the product size was 467 bp. Primer sequences to amplify the insertion event were A-FP together with A-R2: 5’-CAGTAAGCCAAGCGCTACG-3’. In presence of an insertion, a product of 689 bp is amplified. To test the T-DNA insertion, we used primer combination B-FP: 5’-TAAACCAACACTCGGTATGGC-3’and B-RP: 5’-CGGGAGCTTCACACACTTAAG-3’, which amplifies 1001bp in absence of T-DNA insertion. Primer combination B-FP together with LB3 for SAIL lines: 5’-TAGCATCTGAATTTCATAACCAATCTCGATACAC-3’ amplifies ∼500pb in presence of T-DNA insertion.

PCR was run in a total volume of 25uL: 12,5µL EmeraldAmp GT PCR Master Mix (TakaraBio, Code No. RR310A), 0,5 µL of each primer (10µM), 9,5µL milliQ water, 2µL DNA. The following temperature program was used: 96°C for 3 min, 30x (96°C 30 s, 55°C 30 s, 72°C 40 s). PCR products were analyzed by gel electrophoresis on 1% agarose gels.

### Plant transformation

Col-0 WT, *gtr2-1,* and epiRILs573 were transformed with a WT *GTR2* construct in the pFRU vector (Millard et al., 2019) comprising 1972 bp native putative promoter, *GTR2* CDS and 397 bp of the native 3’UTR. For plant transformation, 50 µL electrocompetent *Agrobacterium tumefaciens* cells, C58 (pGV3850), were mixed with 1µL DNA in a 1 mm precooled cuvette and left on ice for 1 min before electroporation at 400 ohm, 2,2 kVm and 25 µF. 1mL YEP medium was added and cells were incubated at 28°C for 2 h. Subsequently, cells were grown on YEP agar plates with antibiotics (50 mg/L spectinomycin, 100 mg/L ampicillin, 69 mg/L rifampicin) at 28°C for two days. 5mL cultures in YEP medium with antibiotics (50 mg/L spectinomycin, 100 mg/L ampicillin, 69 mg/L rifampicin) were started from single colonies and grown overnight at 28°C. Cells were scraped off the plate and suspended in 1 ml 5% (w/v) sucrose solution containing 0,05% (v/v) Silwet L-77. The cell suspension was applied to unopened flower buds. Afterwards, the inflorescences were kept into plastic bags for two days.

### Statistical analysis and visualizations

Outliers were defined by > 2 SD from mean level and were removed from analysis. Data processing was aided by using the package tidyverse, reshape and plyr (Wickham, 2011, 2007; Wickham et al., 2019) Data processing was aided by using the package tidyverse and reshape (Wickham, 2007; Wickham et al., 2019). Statistical analysis was done using R and R studio (R Core Team, 2022; RStudio Team, 2020). Statistical tests were done using the packages rstatix, lme4, lsmeans, agricolae and rcompanion (Bates et al., 2015; Kassambara, 2023a; Lenth, 2016; Mangiafico, 2023). Statistical tests were done using the packages rstatix, lme4, lsmeans and agricolae (Bates et al., 2015; Kassambara, 2023a; Lenth, 2016;Mendiburu, 2023). Plots were made using ggplot2, ggpubr, cowplot, circlize (Gu et al., 2014; Kassambara, 2023b; Wickham, 2016; Wilke, 2020).

### RNA-sequencing

Sequencing was performed on roots of epiRIL573 and WT (epiRIL602 and epiRIL603), 5-6 bioreps per line. RNA was extracted using Spectrum Plant Total RNA kit (Sigma-Aldrich) according to manufacturer’s protocol. Samples were analyzed on a ThermoFisher Nanodrop and concentrations were obtained using ThermoFischer Qubit Fluorometric Quantification with a Qubit RNA HS Assay Kit (Q32855). Libraries were made with the Illumina Neoprep with TruSeq Stranded mRNA Library Prep Kit for NeoPrep (NP-202-1001). Sequencing was performed on an Illumina MiSeq system using the MiSeq Reagent Kit v3 (MS-102-3003). Sequencing was done with Miseq v3 600 cycles (MS-102-3003) and PhiX Control v3 (FC-110-3001) as control. Output files from sequencing were prepared for analysis using Burrows-Wheeler Aligner (BWA) (Li and Durbin, 2009) and Samtools (Li et al., 2009).

Transcriptome analysis was done using R, R Studio (R Core Team, 2022; RStudio Team, 2020) and package DeSeq2 (Love et al., 2014; Zhu et al., 2018) (Supplemental Figure S6). Gene model descriptions were obtained from Arabidopsis.org bulk data retrieval (Berardini et al., 2015). Gene Ontology (GO) enrichment analysis for Biological Processes was carried out using Database for Annotation, Visualization and Integrated Discovery (DAVID, 2023/05/15) (Huang et al., 2009). Of the 439 upregulated DEGs, 398 genes were identified in the enrichment (92.3%). Of the 449 downregulated DEGs, 420 genes were identified in the enrichment (94.4%).

### Global identification of TE insertions

TEs were identified using the program ngs_te_mapper2 (Han et al., 2021). Ngs_te_mapper2 was downloaded using conda (“Anaconda Software Distribution,” 2020). TEs were identified against a TAIR10 genome obtained from the TAIR FTP archive (Berardini et al., 2015). Output was filtered for homozygous TE insertions using a filter on allele frequency > 0,95, uniform 3’ and 5’ support and low reference counts and validated in IGV (Thorvaldsdottir et al., 2013). A global search was done on fastq files of epiRIL573 from ENA project PRJEB5137 (Quadrana et al., 2019). We also tested our data on segregating F2 plants from the epiRIL573 and WT cross. Fastq files were merged into two groups based on having the epiRIL573 phenotype or not.

## Acknowledgement and Funding

We thank Barbara Ann Halkier for providing *gtr2-1* seeds. We thank Marie-Louise Fobian Thomsen and Clarisse Zigue Figueiredo for technical assistance. Financial support for this work was provided by the National Research Foundation DNRF grant 99 (ESTA, CK, PH, MB, DK), The Danish Council for Independent Research, DFF-4181-00077 (ESTA, MB), and Villumfonden, project no. 13169 (ESTA, MB), the U.S. National Science Foundation, Directorate for Biological Sciences, Division of Molecular and Cellular Biosciences, grant no. MCB 1906486 (DJK), the U.S.D.A. National Institute of Food and Agriculture, AFRI project 2019-05709, Hatch project no. CA–D–PLS–7033–H (DJK) and Human Frontier Science Program (RGY0075/2020) (CK).

## Author Contributions

E.S.T.A., D.J.K. and M.B. conceived the project and designed the experiments. E.S.T.A conducted lab work, data analysis, visualization and drafted the manuscript. C.K. generated the complementation constructs and carried out the 8kb promoter complementation. P.H. conducted and analyzed the insect feeding assays. All authors participated in the discussion of the results, and edited and approved the final manuscript. M.B. has ensured that all scientists who have contributed substantially to the conception, design or execution of the work described in the manuscript are included as authors, and that all authors agree to the list of authors and their identified contributions.

